# Genetic diversity and population structure of sweet orange [*Citrus sinensis* (L.) Osbeck] germplasm of India revealed by SSR and InDel markers

**DOI:** 10.1101/2022.01.11.475964

**Authors:** J.Prasanth Tej Kumar, A. Thirugnanavel, Devendra Y. Upadhyay, Snehal A. Kamde, Prafulla R. Jalamkar, Ashutosh A. Murkute

## Abstract

Sweet orange (*Citrus sinensis* (L.) Osbeck) is an important commercial citrus fruit crop, cultivated in India and across the world. In India most of the cultivated sweet orange species were introduced varieties. In this study, we used two molecular markers,SSR and InDels, to understand the genetic diversity and population structure of seventy-two sweet orange genotypes. Genetic parameters consisted of total number of alleles, number of polymorphic alleles (effective alleles); genetic diversity (G.D.), expected heterozygosity (He) and polymorphic information content (PIC) were calculated based on molecular data. Two dendrograms were constructed based on the InDels and SSR. In the both the cases they formed three major clusters showing various degrees of variations with respect to members of the clusters. Population structure analysis revealed presence of two distinct sub populations. Therefore, in order to address various challenges and develop sweet orange varieties with desirable traits, there is a need to broaden the genetic base of sweet orange through intensive collection in the northeastern region. These results of intra-specific genetic variability of the collections will dictate the path for the sweet orange breeding and conservation programs in India.

## Introduction

Citrus (L.) is one of the major fruit crops of the world, belonging to the subfamily Aurantioideae of the family Rutaceae. Citrus is one species which has seamlessly integrated with various cultures across the world with its myriad flavours and grown in different climatic zones like tropical, subtropical and temperate zone, covering more than 140 countries across the world (Kahn et al., 2001). China and Brazil are the major citrus producers with India ranking third (FAO 2018). Citrus in India is grown in 1 million hectares with a total production of 12.55 million tonnes (NH B, 2018).India has an enormous diversity of citrus genetic resources, both cultivated and wild, still untapped.

The genus Citrus (2n=18), comprises of several species like mandarin (*Citrus reticulate*), citron (*Citrus medica*), sweet orange (*Citrus sinensis)*, lime (*Citrus aurantifolia)*, and pummelo (*Citrus maxima*). Sweet orange has major share in the citrus fruits in terms of consumption. In world, sweet orange accounted for approximately 40% of citrus area and 50% of citrus production (FAO,2018). Brazil is the leading country in terms of production of sweet orange followed by China and India (FAO,2018). Sweet orange has the highest consumer preference both as fresh fruit as well as juice. In India, *Citrus sinensis* (L.) Osbeck (sweet orange) occupies the third position after mandarins and limes. In India, the area of sweet orange has been decreased by 3.1% but the production has been increased by 1.75% in the year 2018, indicated the increase in the productivity of sweet orange. In India, presently sweet orange is grown on an area of 0.185 million ha with total production of 3.266 million metric tonnes (NHB, 2018). Sweet oranges are classified into four groups: Common (round oranges), low acidity, pigmented (blood) and navel oranges (Hodgson, 1967; Davies and Albrigo, 1994).

Sweet orange is the first citrus species to be sequenced (Xu et al., 2013) and established as a hybrid between pummelo (*Citrus maxima*) and mandarin (*Citrus reticulata*) (Wu et al., 2018). To understand complexities ofcitrus taxonomy, several phylogenetic studies were done based on morphological and physiological traits (Luro et al., 2011, Gupta et al., 2018), DNA markers (POLAT et al., 2015, Lamine et al., 2015), chrloplast genomes(Carbonell-Caballero et al., 2015), and whole genomes (Wu et al., 2014, Wu et al., 2018,) of several citrus species including sweet orange. Detailed work has been carried out with sweet orange germplasm across the world (Xu et al., 2013, Caruso et al., 2018, Abouzari et al., 2020). But In India, we have barely scratched the tip of the iceberg. Genetic diversity studies in sweet orange have been done in India using RAPD markers (Malik et al., 2012; Sankar et al., 2014).The same genetic materials are often given different names resulting in duplicates when germplasm are assembled from such localities (Teshome et al., 1997). This results in underestimation or overestimation of the actual genetic diversity present in the population (Appa rao et al., 2002).

The identification of markers like SSR, SNP and InDels became more sophisticated with the low cost next-generation sequencing (NGS) technologies (Taheri et al., 2018, Le Nguyen et al., 2019). Simple sequence repeats (SSR) are highly polymorphic, co-dominant, wide genomic distribution and chromosome-specific. In citrus, SSR markers were developed from genomic (ref), ESTs (ref), BAC end sequences (ref). InDels (insertion/deletion) are derived from the insertion of transposable elements, slippage in simple sequence replication or unequal crossover events (Britten et al. 2003). InDels have simple amplicon lengths, high transfer-ability, absence of stutter, less mutation rates and abundant distribution across the genome (Ollitrault et al. 2012, Fang et al., 2018).InDel markers were identified in various citrus species (Ollitrault et al. 2012, Fang et al., 2018, Noda et al., 2020).

In this study, we have taken 72 sweet orange accessions for molecular data analysis using SSR and InDel markers. The objective is to elucidate the phylogenetic relationships, and genetic variability in germplasmwhich can leadto determination of genetic relationships,developing a core collection, designing breeding programs, and validation of new cultivars.

## Materials and methods

### Plant material and collection

A total of 72 cultivars of *Citrus sinensis* (Table 1) were collected from the national active germplasm site of ICAR-Central Citrus Research Institute, Nagpur for molecular studies. Fresh leaves were collected from the selected plant of each cultivar for DNA extraction.

### DNA Extraction

Total genomic DNA was extracted from all the 72 cultivars of sweet orange by using cetyl tri-methyl ammonium bromide (CTAB) method by Doyle and Doyle (1990) with some modifications. Quantitation and purity check of isolated DNA was done using spectrophotometer (BioTek EPOCH|2 microplate reader). DNA was extracted from young leaves using a standard CTAB protocol and the samples were diluted with with 1XTE to 50 ng/μl for PCR reactions.

### PCR Amplification and visualization

A total of 90 SSR were selected based on the chromosome location (Fig 1). These primers were initially screened in 23 citrus accessions comprising of sweet orange, pummelo, mandarin, lime and lemons. Among the 91 primers screened 46 were identified as polymoyphic (unpublished data). Only those 46 SSR primers along with 45 InDel primers selected based on the chromosome lacation were screened in sweet orange accessions. Amplification reaction were performed in the volume of 20μl containing 2μl 10X assay buffer, 1.6 μl of dNTP, 1μl of Taq DNA polymerase (1u/μl), 1μl of DNA primer and 2 μl of template DNA (50 ng/μl). The PCR amplification was performed in BIO-RAD T100 thermal cycler. The PCR amplification conditions were as follows: initial denaturation at 96ºC for 5 minute followed by 45 cycles of denaturation for 30 sec. at 96ºC, annealing at 58ºC (for SSR) and 55ºC (for InDel) for 40 sec. and extension at 72ºC for 1minute followed by final extension at 72ºC for 10 minutes. The amplified product were analysed by electrophoresis (70V for 3 hours) in 4% agarose gel containing ethidium bromide. The gel image was visualized using UV Gel documentation system.

**Fig 1.**
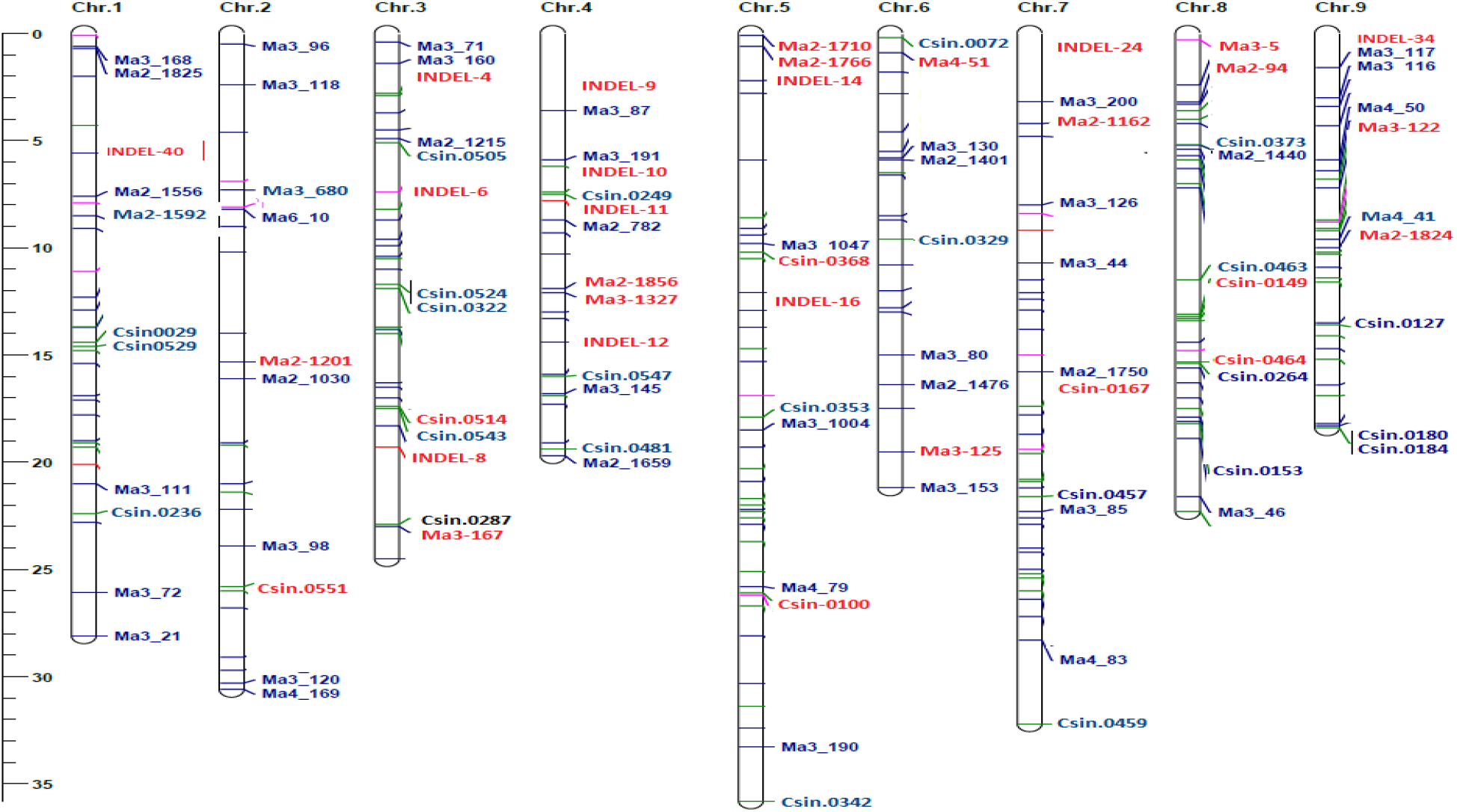
Chromosomal position of SSR and InDels.

### Data Analysis

The alleles of each primer locus were scored (bp) with reference to a known size standard followed by transformation to binary codes as presence (1) or absence (0) of the respective fragment size. A 100 bp DNA ladder (Genexy) was used as standard for scoring. The statistics such as number of alleles per locus, polymorphism % and genetic diversity of each primer were calculated. The polymorphic information content (PIC) and heterozygosity were also calculated for each primer by using the method described by Chesnokov and Artemyeva (2015). The genetic dissimilarity matrix was calculated using simple matching dissimilarity index among pairs of accessions. An UnWeighted neighbor-joining tree was constructed from calculated dissimilarity matrix to depict the genetic relationship among the individuals using the DARwin software version 6 (Perrier and Jacquemoud-Collet, 2006) and the firmness and tenacity of branches was evaluated using 1000 bootstraps.

We used the STRUCTURE software version 2.3.4 (Pritchard et al., 2000) for the determination of the most probable number of clusters for population structure (K value). Using the admixture model, eight simulations were performed for each inferred K value, with a running length composed of 300,000 burning periods and 50,000 Markov chain Monte Carlo (MCMC) iterations to allocate accessions to different populations. The output from this analysis was then used as input in the Structure HARVESTER online program to determine the optimal K value (Earl et al.,2012). the software is based on the ΔK method(Evanno et al.2005).

## Results

### Genetic diversity analysis

In present study, 72 sweet orange germplasms were analysed using 91 molecular markers (45 InDel + 46 SSR), out of which a total of 34 primers [14 InDel (31.11%) and 20 SSR (43.48%)] resulted into 51 polymorphic amplicons (26 InDel + 25 SSR), while 57 [31 InDel (68.89%) + 26 SSR (56.52%)] markers amplified monomorphic products. Allele number ranged from 1-2 with 1.86 and 1.25 average number of alleles per marker of InDel and SSR markers respectively. All the selected InDel and SSR primers have shown 100% polymorphism except IND-chr4-24656 (50% polymorphism). The PIC value depending on allelic frequency in the population (Botstein et al. 1980) ranged from 0.04 (IND-chr4-41717) to 0.48 (IND-chr9-3837) and 0.08 (Ma2-1766) to 0.50 (Csin-0464) with an average value of 0.26 and 0.38 in InDel and SSR markers respectively. A lesser PIC value observed in the present study inferred the presence of low genetic diversity in *Citrus sinensis* germplasms. In the present study, 7 InDel and 12 SSR markers were found to have PIC values more than the average PIC values in respective markers, which shows that these markers are informative and can be useful for trait mapping and tagging studies in Sweet orange germplasm. The heterozygosity ranged from 0.08 (IND-chr3-9113) to 0.91 (IND-chr3-78081) and 0.08 (Ma2-1766) to 1.00 (Csin-0167) with an average value of 0.50 and 0.61 per InDel and SSR markers respectively. Genetic diversity values were observed in the range of 0.04 (IND-chr4-41717) to 0.49 (IND-chr9-3837) and 0.08 (Ma2-1766) to 0.51 (Csin-0464) with an average value of 0.26 and 0.39 per InDel and SSR markers respectively.

### Cluster analysis by Indels

Cluster analysis based on InDel markers data and neighbour joining tree divided all the 72 accessions into three main clusters (Fig. 1). Among all the clusters, cluster I was the largest comprising of 43 accessions, cluster II comprised of 25 accessions and cluster III was the smallest comprising of only 4 accessions. The first cluster is again subdivided into two major clusters. The IA cluster contains 30 accessions among them M53 and M37; M69 and M52 have higher similarity. The IB contains 13 accessions with no duplicates. The northeastern collections like Soh Bittara, Soh nairiang, Soh Khylla and Tasi found in this cluster. The II cluster contains 25 accessions. Highest number of duplications were found in this cluster. In the III cluster only four accessions were found.

### Cluster analysis by SSR

Cluster analysis based on SSR data and neighbour joining tree divided all the 72 accessions into three main clusters (Fig. 2). There are several subclusters among each cluster. The cluster I contains 36 accessions. The cluster II contains 30 accessions and the cluster III contains 6 accessions

**Fig 2.**
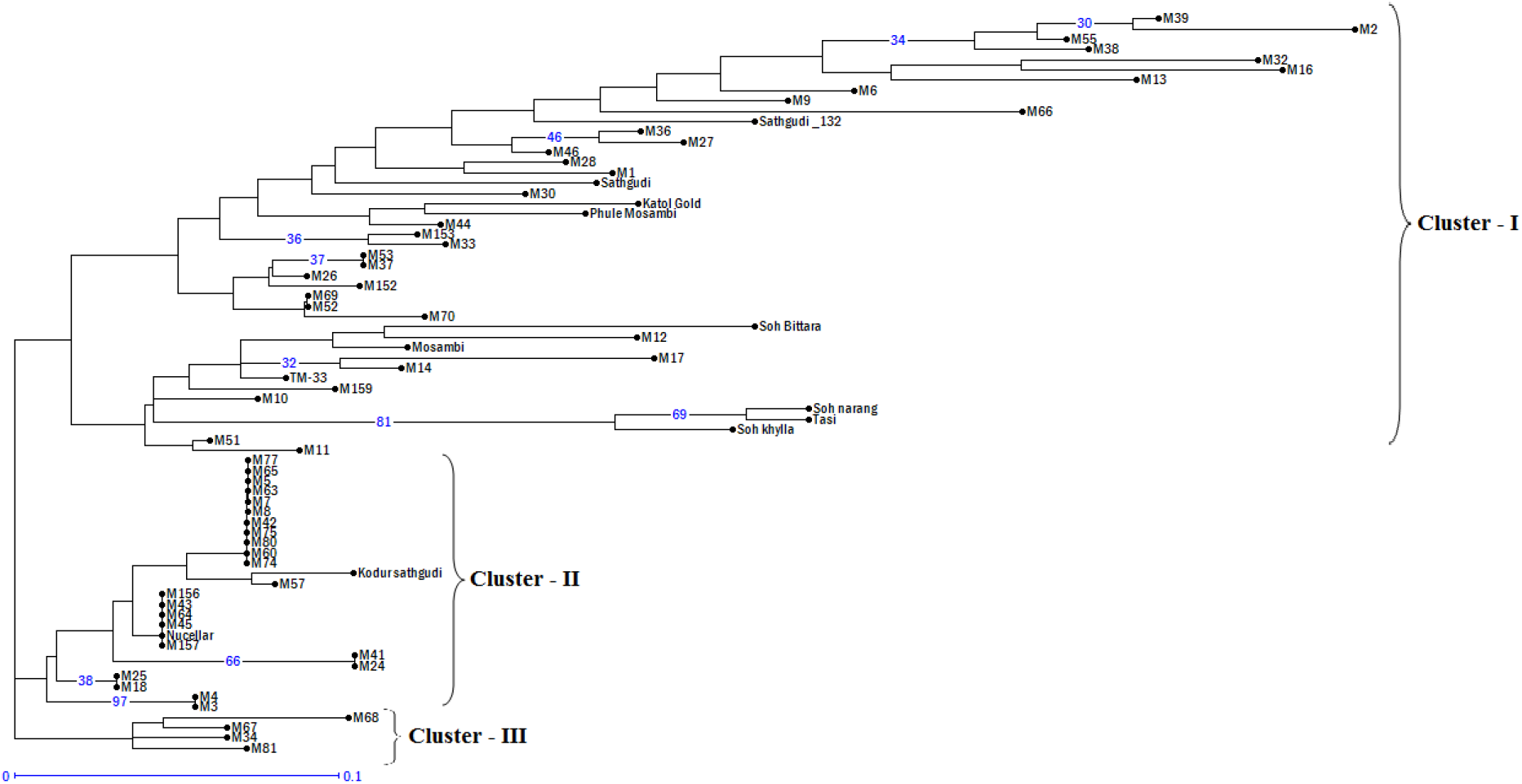
Dendrogram of 72 sweet orange accessions based on InDels.

**Fig 3.**
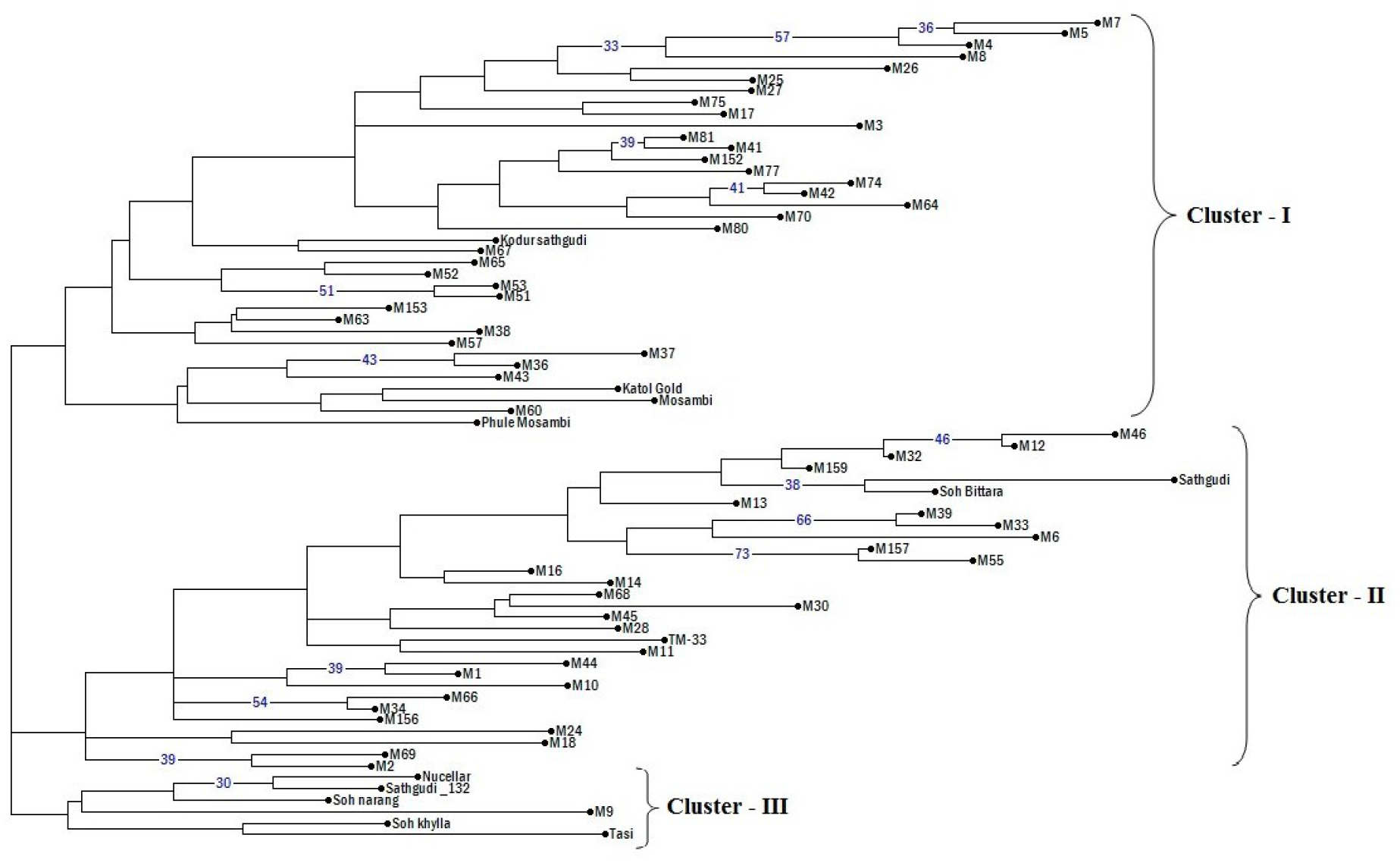
Dendrogram of 72 sweet orange accessions based on SSR.

### Population structure analysis

The calculated membership fractions of 72 clones for different k values were varying from 1 to 6. The log likelihood determined by STRUCTURE revealed the optimal value 2 (K = 2). Correspondingly, the highest of adhoc measure (ΔK) analysis (Evanno et al., 2005) gave a higher likelihood value at K = 2 (Fig 4), which demonstrated the occurrence of 2 sub-populations in entire accession set used in study (Fig 5).

**Fig. 4:**
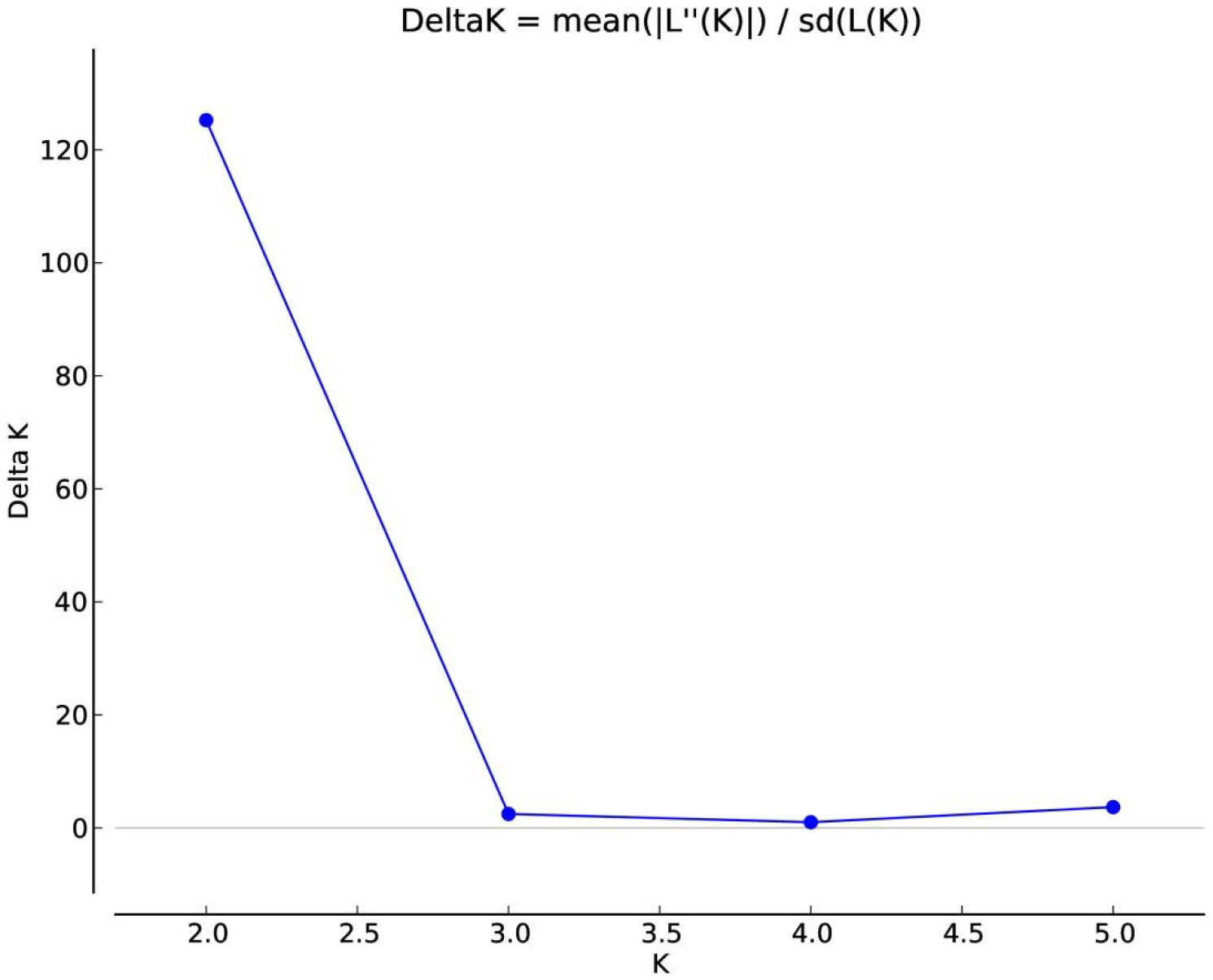
Estimation of population using LnP(D) derived ΔK with K ranged from 1 to 6 with SSR and InDels.

**Fig 5.**
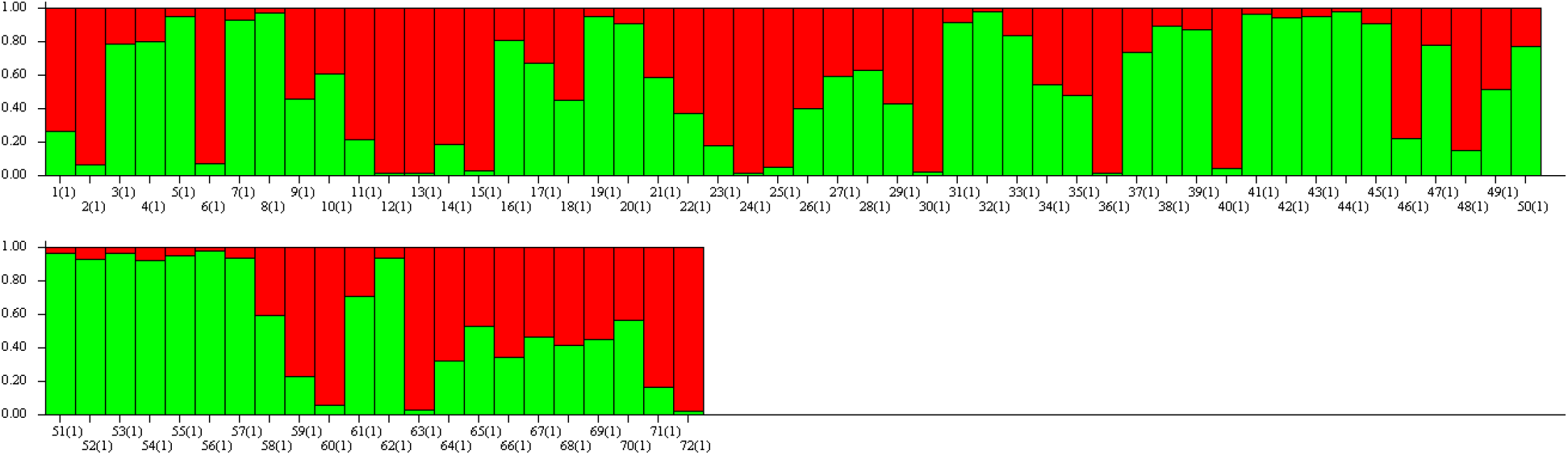
Population structure of 72 sweet orange accessions based on SSR and Indel markers (K = 2) and graph of estimated membership fraction for K = 2.

## Discussion

Sweet orange is known for its taste and commercially most important crop across the world. The fruit mostly consumed in the form of juice and most of the Indian juice consumption is coming from the foreign countries. Indian varieties are not tenable for the processing industries (Ref). Despite being the one of the origin of the centers we are yet to tap the full potential of the fruit. In India, sweet orange occupies second place after the mandarins. The main reason for this may be sweet orange is irrigation intensive and less market value. The genotypic data of the sweet orange accessions was used for the analyzing genetic diversity and population genetics, which might shape up future breeding efforts in *Citrus sinensis*

In this study we have assessed the genetic diversity based on the InDel and SSR. Most of the accessions taken in this study were either clonal selection or bud mutations. Some of the cultivars collected from the northeastern region. There are clear differences in the clusters formed with respect to InDel and SSR. The most prominent was the presence of the duplicates in the Indel based clusters and there were none in SSR based clusters. The soh khylla, soh niairang and Tasi have grouped in the same clusters as expected but soh bittara has moved away to different cluster with SSR data. Cluster analysis clearly separates the Indian cultivars like Tasi, soh Khyllah and soh nairiang in both the cases, except soh bittara. The Katol gold, mosambi and Phule mosambi have grouped similarly in the both the clusters. Katol gold another sweet orange variety getting prominence in recent years. The kodur sathgudi which was supposed to be the clonal selection of sathgudi is grouping in different cluster with respect to sathgudi. This can be further verified by using more number of markers. Most of our ruling varieties are the exotic species like mosambi and sathgudi.

Most of the sweet oranges are diploids with a comparatively small genome size of about 367 Mb (Arumuganathan and Earle, 1991, *Latest draft genome reference*).Sweet oranges usually show low level of genetic diversity (Novelli et al., 2006; Jannati et al. 2009; Polat, 2014). After coming through by introduction, most Indian sweet orange accessions originated via mutations. Besides there are some Indian origin collections which show clear separation from the others. The level of polymorphism is very less may be due to the narrow genetic basis and somatic mutations contributes for most of the variation. The genetic diversity of *C.sinensis* is reducing due to selection and introduction of genotypes suitable for intensive horticulture (**Agaro, 2000**). The sweet orange genetic resources in India apparently have been subjected to human selection for centuries is creating a genetic bottleneck

Sweet orange is a result of the natural cross between pummelo and mandarin. The population structure showing two sub populations might belong to mandarin and pummelo .Several online based citrus genetic resources databases were coming up and data from India will help to speed up the breeding processes and to protect the species from extinction (Biswas et al., 2020). The molecular characterization combined with morphological characters will help to assess the genetic diversity. In the long run helps for the genetic improvement of the sweet orange.

## Conclusions

In the present study, the genetic diversity and population structure of sweet orange germplasm resources were analyzed using SSR and InDels. The results showed they have abundant genetic diversity. However, the results vary with respect to SSR and Indels. With area increasing in the sweet orange, its time we work towards the coloured sweet oranges and varieties suitable for processing.

