## Supplementary material for "Genetic diversity and population structure of sweet orange [*Citrus sinensis* (L.) Osbeck] germplasm of India revealed by SSR and InDel markers": Tables

**Table 1 :Details of sweet orange accessions**

| Code no. | Name of the germplasm | IC/EC no. | Source of origin |
|---|---|---|---|
| 1 | M1 | IC-311476 | Jadgaon, Aurangabad, Maharashtra |
| 2 | M2 | IC-311478 | Dawarwadi, Aurangabad, Maharashtra |
| 3 | M3 | IC-311479 | Paithan, Aurangabad, Maharashtra |
| 4 | M4 | IC-311480 | Paithan, Aurangabad, Maharashtra |
| 5 | M5 | IC-311481 | Paithan, Aurangabad, Maharashtra |
| 6 | M6 | IC-311482 | Paithan, Aurangabad, Maharashtra |
| 7 | M7 | IC-311483 | Paithan, Aurangabad, Maharashtra |
| 8 | M8 | IC-311484 | Dawarwadi, Jalna, Maharashtra |
| 9 | M9 | IC-311485 | Talegaon, Jalna, Maharashtra |
| 10 | M10 | Obtained through clonal selection | ICAR-CCRI, Nagpur, India |
| 11 | M11 | Obtained through clonal selection | ICAR-CCRI, Nagpur, India |
| 12 | M12 | Obtained through clonal selection | ICAR-CCRI, Nagpur, India |
| 13 | M13 | IC-311489 | Guha, Ahamadnagar, Maharashtra |
| 14 | M14 | Obtained through clonal selection | ICAR-CCRI, Nagpur, India |
| 15 | M16 | IC-311490 | Adhawadi, Ahamadnagar, Maharashtra, |
| 16 | M17 | IC-311491 | Kolgaon, Ahamadnagar, Maharashtra |
| 17 | M18 | Obtained through clonal selection | ICAR-CCRI, Nagpur, India |
| 18 | M24 | Obtained through clonal selection | ICAR-CCRI, Nagpur, India |
| 19 | M25 | IC-311493 | Kolgaon, Ahamadnagar, Maharashtra |
| 20 | M26 | Obtained through clonal selection | ICAR-CCRI, Nagpur, India |
| 21 | M27 | Obtained through clonal selection | ICAR-CCRI, Nagpur, India |
| 22 | M28 | Obtained through clonal selection | ICAR-CCRI, Nagpur, India |
| 23 | M30 | Obtained through clonal selection | ICAR-CCRI, Nagpur, India |
| 24 | M32 | Obtained through clonal selection | ICAR-CCRI, Nagpur, India |
| 25 | M33 | Obtained through clonal selection | ICAR-CCRI, Nagpur, India |
| 26 | M34 | Obtained through clonal selection | ICAR-CCRI, Nagpur, India |

| | | | |
|---|---|---|---|
| 27 | M36 | Obtained through clonal selection | ICAR-CCRI, Nagpur, India |
| 28 | M37 | Obtained through clonal selection | ICAR-CCRI, Nagpur, India |
| 29 | M38 | IC-311494 | Kolgaon, Ahamadnagar, Maharashtra |
| 30 | M39 | Obtained through clonal selection | ICAR-CCRI, Nagpur, India |
| 31 | M41 | IC-311495 | Kolgaon, Ahamadnagar, Maharashtra |
| 32 | M42 | Obtained through clonal selection | ICAR-CCRI, Nagpur, India |
| 33 | M43 | Obtained through clonal selection | ICAR-CCRI, Nagpur, India |
| 34 | M44 | Obtained through clonal selection | ICAR-CCRI, Nagpur, India |
| 35 | M45 | IC-311498 | Talegaon, Pune, Maharashtra |
| 36 | M46 | IC-311501 | Shokrapur, Pune, Maharashtra |
| 37 | M51 | Obtained through clonal selection | ICAR-CCRI, Nagpur, India |
| 38 | M52 | Obtained through clonal selection | ICAR-CCRI, Nagpur, India |
| 39 | M53 | Obtained through clonal selection | ICAR-CCRI, Nagpur, India |
| 40 | M55 | Obtained through clonal selection | ICAR-CCRI, Nagpur, India |
| 41 | M57 | Obtained through clonal selection | ICAR-CCRI, Nagpur, India |
| 42 | M60 | IC-311502 | Usthaldumali, Ahamadnagar, Maharashtra |
| 43 | M63 | IC-311503 | Usthaldumali, Ahamadnagar, Maharashtra |
| 44 | M64 | IC-311504 | Kaygaon, Aurangabad, Maharashtra |
| 45 | M65 | Obtained through clonal selection | ICAR-CCRI, Nagpur, India |
| 46 | M66 | Obtained through clonal selection | ICAR-CCRI, Nagpur, India |
| 47 | M67 | IC-311505 | Kaygaon, Aurangabad, Maharashtra |
| 48 | M68 | Obtained through clonal selection | ICAR-CCRI, Nagpur, India |
| 49 | M69 | IC-322089 | Baruasagar, Jhansi, U.P. |
| 50 | M70 | Obtained through clonal selection | ICAR-CCRI, Nagpur, India |
| 51 | M74 | IC-322097 | Baruasagar, Jhansi, U.P. |
| 52 | M75 | Obtained through clonal selection | ICAR-CCRI, Nagpur, India |
| 53 | M77 | IC-322247 | Mahatargaon, Hingoli, Maharashtra |
| 54 | M80 | Obtained through clonal selection | ICAR-CCRI, Nagpur, India |
| 55 | M81 | Obtained through clonal selection | ICAR-CCRI, Nagpur, India |

| | | | |
|---|---|---|---|
| 56 | M152 | Obtained through clonal selection | ICAR-CCRI, Nagpur, India |
| 57 | M153 | Obtained through clonal selection | ICAR-CCRI, Nagpur, India |
| 58 | M156 | Obtained through clonal selection | ICAR-CCRI, Nagpur, India |
| 59 | M157 | Obtained through clonal selection | ICAR-CCRI, Nagpur, India |
| 60 | M159 | Obtained through clonal selection | ICAR-CCRI, Nagpur, India |
| 61 | Phule Mosambi | Obtained from State Agricultutral University | Mahatma Phule Krishi Vidyapeeth (MPKV), Rahuri, Maharashtra |
| 62 | Kodur sathgudi | Obtained from State Agricultutral University | Citrus Research Station, Tirupati (DR. YSR Horticultural University), Andhra Pradesh |
| 63 | Soh Bittara | IC- 344929 | Barapani, Ribhoi, Meghalaya |
| 64 | Mosambi | Obtained through clonal selection | ICAR-CCRI, Nagpur, India |
| 65 | Katol Gold | Obtained from State Agricultutral University | Dr. Panjabrao Deshmukh Krishi Vidyapeeth, PDKV, Akola, Maharashtra, |
| 66 | Sathgudi _132 | Obtained through clonal selection | ICAR-CCRI, Nagpur, India |
| 67 | Tasi | IC- 344927 / IC- 346977 | Arunachal Pradesh |
| 68 | Soh khylla | IC-344935 | Umroi, Ribhohi, Meghalaya |
| 69 | Soh nairiange | IC- 346992 / IC-344928 | Basar, West Siang, Arunachal Pradesh / Barapani, Ribhoi, Meghalaya |
| 70 | Nucellar | Obtained through clonal selection | ICAR-CCRI, Nagpur, India |
| 71 | TM-33 | Obtained through clonal selection | ICAR-CCRI, Nagpur, India |
| 72 | Sathgudi | IC-322250 | Limbgaon, Nanded, Maharashtra |

**Table 2 . Polymorphic analysis of InDel primers used in Sweet Orange**

| Sr. No. | InDel Primer | Total no. amplicon | Poly. alleles | Polymorphism % | PIC | G.D. | Heterozygosity |
|---|---|---|---|---|---|---|---|
| 1 | IND-chr3-9113 | 2 | 1 | 50 | 0.08 | 0.08 | 0.08 |
| 2 | IND-chr3-78081 | 1 | 1 | 100 | 0.41 | 0.42 | 0.91 |
| 3 | IND-chr3-100795 | 2 | 2 | 100 | 0.09 | 0.09 | 0.55 |
| 4 | IND-chr4-1894 | 2 | 2 | 100 | 0.33 | 0.33 | 0.37 |
| 5 | IND-chr4-24656 | 2 | 1 | 50 | 0.05 | 0.05 | 0.50 |
| 6 | IND-chr4-41717 | 2 | 2 | 100 | 0.04 | 0.04 | 0.50 |
| 7 | IND-chr4-67425 | 2 | 2 | 100 | 0.39 | 0.39 | 0.46 |
| 8 | IND-chr5-7677 | 2 | 2 | 100 | 0.42 | 0.42 | 0.51 |
| 9 | IND-chr5-24964 | 2 | 2 | 100 | 0.12 | 0.12 | 0.58 |
| 10 | IND-chr6-69785 | 1 | 1 | 100 | 0.44 | 0.45 | 0.56 |
| 11 | IND-chr7-2104 | 2 | 2 | 100 | 0.20 | 0.20 | 0.21 |
| 12 | IND-chr9-3837 | 2 | 2 | 100 | 0.48 | 0.49 | 0.64 |
| 13 | IND-chr1-10341 | 2 | 2 | 100 | 0.38 | 0.38 | 0.45 |
| 14 | IND-chr1-43527 | 2 | 2 | 100 | 0.20 | 0.21 | 0.65 |
| | **Mean** | 1.86 | 1.79 | 96.43 | 0.26 | 0.26 | 0.50 |

**Note: PIC – Polymorphic Information Content and G.D. – Genetic Diversity.**

**Table 3. Polymorphic analysis of SSR primers used in Sweet Orange**

| Sr. No. | SSR Primer | Total no. amplicon | Poly. alleles | Polymorphism % | PIC | G.D. | Heterozygosity |
|---|---|---|---|---|---|---|---|
| 1 | Csin-0551 | 1 | 1 | 100.0 | 0.42 | 0.43 | 0.52 |
| 2 | Ma3-167 | 1 | 1 | 100.0 | 0.10 | 0.11 | 0.11 |
| 3 | Ma2-1766 | 1 | 1 | 100.0 | 0.08 | 0.08 | 0.08 |
| 4 | Csin-0100 | 2 | 2 | 100.0 | 0.35 | 0.35 | 0.88 |
| 5 | Ma4-51 | 1 | 1 | 100.0 | 0.48 | 0.48 | 0.63 |
| 6 | Csin-0167 | 1 | 1 | 100.0 | 0.42 | 0.43 | 0.91 |
| 7 | Ma2-94 | 2 | 2 | 100.0 | 0.41 | 0.42 | 0.91 |
| 8 | Ma2-1824 | 1 | 1 | 100.0 | 0.49 | 0.50 | 0.68 |
| 9 | Csin-0368 | 1 | 1 | 100.0 | 0.47 | 0.48 | 0.61 |
| 10 | Ma3-1327 | 1 | 1 | 100.0 | 0.35 | 0.35 | 0.40 |
| 11 | Ma3-125 | 2 | 2 | 100.0 | 0.46 | 0.47 | 0.85 |
| 12 | Ma2-1162 | 1 | 1 | 100.0 | 0.35 | 0.35 | 0.40 |
| 13 | Csin-0149 | 1 | 1 | 100.0 | 0.48 | 0.49 | 0.64 |
| 14 | Ma3-122 | 1 | 1 | 100.0 | 0.31 | 0.32 | 0.35 |
| 15 | Csin-0514 | 2 | 2 | 100.0 | 0.43 | 0.44 | 0.54 |
| 16 | Ma3-5 | 2 | 2 | 100.0 | 0.13 | 0.13 | 1.00 |
| 17 | Ma2-1201 | 1 | 1 | 100.0 | 0.49 | 0.50 | 0.68 |
| 18 | Ma2-1710 | 1 | 1 | 100.0 | 0.38 | 0.38 | 0.44 |
| 19 | Csin-0464 | 1 | 1 | 100.0 | 0.50 | 0.51 | 0.72 |
| 20 | Ma2-1856 | 1 | 1 | 100.0 | 0.49 | 0.50 | 0.80 |
|  | **Mean** | 1.25 | 1.25 | 100.00 | 0.38 | 0.39 | 0.61 |
